# Gut inflammation promotes adverse food reactions by disrupting the microbial metabolism of food triggers

**DOI:** 10.64898/2026.02.05.703147

**Authors:** Bruna B. Da Luz, Liam E. Rondeau, Rebecca Dang, Dana Coppens, Dominika Boron, Pranshu Muppidi, Jessica Linton, Fernando A. Vicentini, John K. Marshall, Giada De Palma, Premysl Bercik, Neeraj Narula, Alberto Caminero

## Abstract

**Background & Aims:** Food-related adverse reactions are frequently reported by patients with inflammatory bowel disease (IBD), but the underlying mechanisms are poorly understood. We investigated how intestinal inflammation and the microbiota contribute to the development of adverse food reactions.

**Methods:** We sensitized mice to different foods (dairy and gluten) after intestinal inflammation (chemically- and hapten-induced models), and then re-exposure to the sensitized foods through diet enrichment. To study whether inflamed microbiota facilitates adverse food reactions, we employed gnotobiotic models and bacterial supplementation experiments. We assessed markers of intestinal inflammation and sensitization, clinical responses, RNA transcripts, and microbiota composition and function. In a translational approach, we recruited IBD patients in remission and healthy controls, recorded self-reported food intolerances and clinical responses to triggering foods, and feces for gut microbiota analyses were collected.

**Results:** Intestinal inflammation facilitates food sensitization by disrupting microbial antigen metabolism, recruiting mast cells to the colon, and promoting mucosal IgE production. In gnotobiotic models, inflammation-driven depletion of colonic bacteria involved in food digestion contributed to food sensitization. Upon re-exposure to triggering foods, sensitized mice experienced visceral pain and low-grade inflammation through mast-cell mediated mechanisms, which also worsened experimental colitis. Supplementation with depleted bacteria or treatment with mast cell stabilizers attenuated food-driven responses. In IBD patients, self-reported food intolerances were common and associated with microbial disruption and depletion of food-metabolizing bacteria.

**Conclusion:** Microbial metabolism of foods is disrupted after intestinal inflammation. This facilitates food sensitization, through colonic mast cell-mediated immune responses, which may explain the high number of adverse food reactions reported by IBD patients.

**WHAT YOU NEED TO KNOW:** *Background and context:* Patients with inflammatory bowel disease (IBD) frequently report adverse food reactions, but the underlying mechanisms are not well understood.

*New findings:* Intestinal inflammation promotes food sensitization by depleting bacteria that degrade food triggers. IBD patients in remission with food intolerances show reduced microbial diversity and loss of bacteria involved in digesting food triggers.

*Limitations:* We used chemically- and hapten-induced mouse models in this study due to the importance of monitoring inflammation onset. Food-driven immune reactions in the mucosa of IBD patients were not performed.

*Clinical research relevance:* Impaired microbial food metabolism is linked to adverse food reactions in IBD. Microbiome-based therapies, such as probiotics capable of degrading dairy or gluten, should be considered for IBD patients with food intolerances.

*Basic research relevance:* We identified a novel mechanism in which microbial disruption caused by intestinal inflammation leads to adverse food reactions and worsened colitis in preclinical models. Restoring the microbial capacity to digest trigger foods reverses these effects.

*Lay Abstract:* Intestinal inflammation facilitates sensitization to gluten and dairy proteins by depleting microbes that digest them, contributing to the increase in adverse food reactions among IBD patients.

## Introduction

Inflammatory bowel disease (IBD), which includes Crohn’s disease (CD) and ulcerative colitis (UC), is a chronic relapsing condition of increasing prevalence, characterized by episodes of severe inflammation and tissue damage^1^. In addition to pharmacological therapies, which are expensive and limited in long-term efficacy, many patients modify their diets to manage their symptoms^2^. Although scientific evidence supporting targeted dietary interventions in IBD is limited, elimination diets are frequently adopted by patients seeking symptom relief. Approximately two-thirds of patients with IBD report adverse reactions to foods^3,4^, and common food sensitivities such as celiac disease or food allergy, are more prevalent among individuals with IBD^5,6^. The most commonly self-reported food triggers among IBD patients include dairy, wheat, and fiber^7^, but the mechanisms underlying these adverse reactions remain unknown.

There is growing evidence that altered diet-microbiota interactions contribute to the development and course of IBD^8,9^. The intestinal microbiota is considered a secondary metabolic organ, contributing to the digestion of foods that human enzymes cannot process, such as common food triggers and antigens^10,11^. Moreover, the intestinal microbiota has been associated with the development of adverse reactions to foods^12,13^. Several studies have shown that IBD patients present both altered compositional and metabolic function of the gut microbiota^14,15^. Nevertheless, the role of the intestinal microbiota in the development of food-related adverse reactions in IBD is not fully understood.

Here, we show that gut inflammation promotes sensitization to foods, such as dairy and gluten proteins, using two murine models of colitis. Sensitization to dairy and gluten proteins after inflammation resulted in innate immune activation, the generation of mucosal IgE, and mast cell recruitment in the colon. Reintroducing trigger foods resulted in visceral hypersensitivity, low-grade inflammation, and exacerbation of experimental colitis. Inflammation significantly altered the intestinal microbiota, depleting bacteria involved in dairy and gluten metabolism and impairing food antigen processing, which contributed to food sensitization in gnotobiotic mice. Notably, microbiota transfer from inflamed to germ-free mice was sufficient to promote sensitization, while supplementation with antigen-metabolizing bacteria mitigated these effects. Finally, we identified a higher prevalence of self-reported food intolerances in IBD patients, associated with reduced complexity of the fecal microbiota and depletion of food-metabolizing bacterial taxa. Together, these findings reveal a mechanism by which inflammation drives adverse food reactions by disrupting microbial metabolism of dietary antigens and activating colonic mast cell-mediated immune responses.

## Materials and Methods

### Participant Recruitment

A total of 148 participants (58 CD, 46 UC, and 44 non-IBD controls) were recruited from the Gastroenterology clinic at McMaster University between August 2021 and September 2023. Inclusion criteria for IBD participants included a confirmed diagnosis of CD or UC between the ages of 18–75 years old, and clinical and histological remission. Non-IBD controls aged 18–75 and undergoing colonoscopy for colorectal cancer screening with no organic disease or IBD history, were recruited for the study. Exclusion criteria for all participants included inability to provide written consent, concurrent systemic disease, history of active cancer in the past 5 years, life-threatening comorbidities, pregnancy or breastfeeding, recent use of antibiotics (within the past month), and any immunocompromising condition. The study was approved by the Hamilton Integrated Research Ethics Board (HiREB #13741). All participants provided written informed consent before the study, completed questionnaires on food intolerances, symptoms, and demographics, and supplied stool samples for 16S rRNA Illumina sequencing to assess microbial composition. Fecal samples were processed anaerobically and stored at −80°C until analysis.

### Preclinical experimental design

To study the involvement of gut inflammation in food sensitization, 6-to-8-week-old specific pathogen-free (SPF) C57BL/6 mice (Taconic) were used. Colitis was induced using 3% dextran sulfate sodium (DSS) or 2% 2,4,6-trinitrobenzene sulfonic acid (TNBS), followed by sensitization to food proteins (dairy and gluten) using cholera toxin (Cedarlane). Mice were then fed isocaloric customized diets containing or excluding dairy or gluten for one week, with or without a second cycle of 1.5% DSS during the dietary intervention.

In a subset of mice, cromolyn sodium (100mg/kg) was administered daily via intraperitoneal injection during dietary interventions to evaluate the role of mast cells in inflammation-induced food reactions. Another subset received daily oral supplementation of *Clostridium sporogenes* (10⁹ CFU/mouse), isolated from healthy volunteers ^16^, to evaluate the contribution of dairy protein-degrading bacteria in mitigating adverse food reactions.

For gnotobiotic experiments, germ-free C57BL/6 mice were generated through two-stage embryo transfer and bred under gnotobiotic conditions in the Axenic Gnotobiotic Unit (AGU) at McMaster University. To study the role of inflammation-driven microbiota alterations in food sensitization, germ-free C57BL/6 were colonized with SPF Non-DSS or post-DSS microbiota via oral gavage with 200 uL of microbial suspension slurries. Microbial suspensions were prepared by diluting fecal content 1:10 in sterile phosphatase-buffer saline. Mice were then sensitized to dairy proteins and fed the dairy-protein diet. All experiments were approved by the McMaster University Animal Care Committee and conducted in accordance with the Animal Utilization Protocol (#22-26), and ARRIVE guidelines 2.0.

### Intestinal inflammation models

Intestinal injury was induced by DSS or TNBS. In DSS-induced inflammation, C57BL/6 mice were exposed to 3% DSS (36,000–50,000 MW; MP Biomedicals LLC) in drinking water for 5 days, followed by a recovery period with normal drinking water. Controls mice were exposed to water instead of DSS. In TNBS-induced inflammation, mice were anesthetized with isoflurane (4%), and TNBS (2% diluted in 50% ethanol) was administered by rectal instillation using a polyethylene catheter. Control mice underwent the same procedure but were administered vehicle (50% ethanol) instead of TNBS. Four hours later, the animals were given free access to food and water^17^. Colitis severity was evaluated at endpoint by measuring colon length, body-weight loss, a disease activity index (DAI). Stool consistency and blood were each scored on a scale of 0–3, as previously described, then summed for a DAI (Supplementary Materials). Histologic damage was assessed in haematoxylin and eosin-stained distal colon cross sections. Polymorphonuclear cells (PMNs) were quantified in colon cross sections by counting the number of PMNs and measuring tissue area in 2-3 cross sections per mouse. For each section, the number of PMNs per µm^2^ was calculated and these values were averaged to obtain a single mean value per mouse. All quantifications were performed in a blinded manner.

### Oral sensitization to dairy and gluten proteins

To break oral tolerance to food, mice were intragastrically administered 6 mg of casein plus 6 mg of whey protein (Sigma-Aldrich), 6 mg of gluten (Sigma-Aldrich), or vehicle, each combined with 10 µg of cholera toxin (Cedarlane) on days 4, 8, 12, and 16 following induced inflammation by DSS or TNBS. Classic markers of food sensitization (mucosal IgE and colonic mast cells) and food-driven immune reactions (mucosal mast cell protease 1; mMCPT-1) were analyzed (Supplementary Materials).

### Diet interventions

Four isocaloric, customized diets were formulated in collaboration with Envigo: a wheat containing diet (TD.2919); a dairy-protein diet using casein and whey in a 4:1 ratio, reflecting the natural composition of milk (TD.220453); a wheat-free diet (TD.05620); and a wheat-containing diet (TD.200056). All diets present the same amount of protein (14%).

### Fecal and bacterial capacity to degrade food triggers

Bacteria isolated from human feces (Caminero Lab collection^16^) and mouse feces were tested for their capacity to degrade dairy proteins. Briefly, bacterial strains, or feces diluted 1/10 (w/v) in PBS, were incubated in liquid BHI, supplemented with casein (Sigma - 1%) and whey (Sigma - 1%). After 48 h of incubation in anaerobic conditions, the amount of Bos d 5 and Bos d 11 (β-casein and β-lactalbumin proteins) were quantified by ELISA (Indoor Biotechnology). Degradation capacity was displayed as a semi-quantitative measure by comparing the initial concentration of Bos d 5 and Bos d 11 in the non-growth negative control to the remaining concentration in the experimental samples. The data are represented as heatmaps where red=100% degradation, and white=0% degradation.

### Microbiota analysis

Genomic DNA was extracted from mice and human feces, and the variable V3-V4 region of 16S rRNA were amplified with polymerase chain reaction (PCR) using Taq polymerase (Life Technologies, Carlsbad, CA). Forward barcoded primers targeting the V3 region (v3f_341f-CCTACGGGNGGCWGCAG) and reverse primers targeting the V4 region (v4r_806r-GGACTACNVGGGTWTCTAAT) were used. Forward primers included six-base pair barcodes to allow multiplexing samples. Purified PCR products were sequenced using the Illumina MiSeq platform by the McMaster Genomics facility. Primers were trimmed from the obtained sequences with Cutadapt software^19,20^, and processed with Divisive Amplicon Denoising Algorithm 2 (DADA2; version 1.14.0) using the trained SILVA reference database (version 138.1)^21,22^. For mouse microbiota, a total of 5,612,228 reads were obtained with a minimum of 12,875 reads and maximum of 97,477 reads, with an average of 62,258 read per mouse sample. For human microbiota, a total of 5,292,618 reads, with a minimum of 4,821 reads and maximum of 123,105 reads, with an average of 41,674 reads per individual sample. Gut microbiota α-diversity was measured by Shannon index and β-diversity was calculated using Bray-Curtis dissimilarity measures. Analysis was completed in RStudio software, version 2023.12.0^+^, using Phyloseq package (version 1.46.0).

### Statistical methods

All variables were analyzed using SPSS version 26 (SPSS Inc., USA) and GraphPad Prism 9 (GraphPad Software, USA). Statistical analysis was performed using Kruskal–Wallis followed by Dunn’s test for non-parametric data, one-way or two-way ANOVA followed by Bonferronís test for parametric data or unpaired t test. Normal distribution was determined by Kolmogorov– Smirnov test. A P value < 0.05 was selected to reject the null hypothesis. Details on specific p-values and data presentation are provided in the figure legends.

## Results

### Intestinal inflammation facilitates sensitization to dairy and gluten proteins

To determine how intestinal inflammation facilitates adverse food reactions, we established a mouse model of food protein sensitivity (Figure 1A). C57BL/6 mice were given DSS to induce colitis and sensitized to dairy proteins using cholera toxin or received sham sensitization. After recovery, mice were provided either a dairy-containing or control diet. All DSS-treated mice experienced weight loss and increased disease activity index (DAI) during colitis, which resolved after recovery (12–14 days). However, only mice sensitized during inflammation and subsequently fed a dairy-containing diet exhibited a secondary rise in the DAI during the dietary intervention (Figure 1B). Colonic mucosal inflammation was more evident in DSS-treated mice compared to non-DSS controls (Supplementary Figure 1A-C), but there were no differences in weight loss, colon length, or histologic damage across DSS-treated groups. Importantly, the colonic mucosa of dairy-sensitized mice fed a dairy diet had significantly higher numbers of polymorphonuclear cells (PMNs) and CD3^+^ cells compared to non-sensitized and non-DSS controls (Figure 1C and 1D). To explore immune mechanisms underlying these responses, gene expression in the colonic tissue was evaluated. Nine pro-inflammatory genes were upregulated in sensitized mice on a dairy diet, including Th1-related (*Cxcl9*, *Cxcl10*), inflammatory (*Tnf*, *Il-1a*, *Il-6*), and apoptosis-associated genes (*Arg1*, *C6*) (Figure 1E, Supplementary Figure 1D). Consistent with these findings, increased TUNEL-positive cells confirmed enhanced mucosal apoptosis in sensitized mice fed a dairy diet (Figure 1F). Key mediators of food sensitivity were then assessed. Dairy-fed, sensitized mice exhibited increased leukocyte migration to the abdominal cavity (Supplementary Figure 1E) and elevated colonic IgE compared to non-DSS and non-sensitized controls, which suggests immune sensitization to dairy is increased after inflammation (Figure 1G). Finally, because abdominal pain is the most common self-reported dietary-driven symptom in IBD patients^3^, we investigated visceral sensitivity in mice exposed to DSS and sensitized to dairy or sham, then fed a dairy diet. Sensitized mice displayed heightened visceromotor responses to colorectal distension, indicating sensitized-mediated visceral hypersensitivity (Figure 1H).

**Figure 1.**
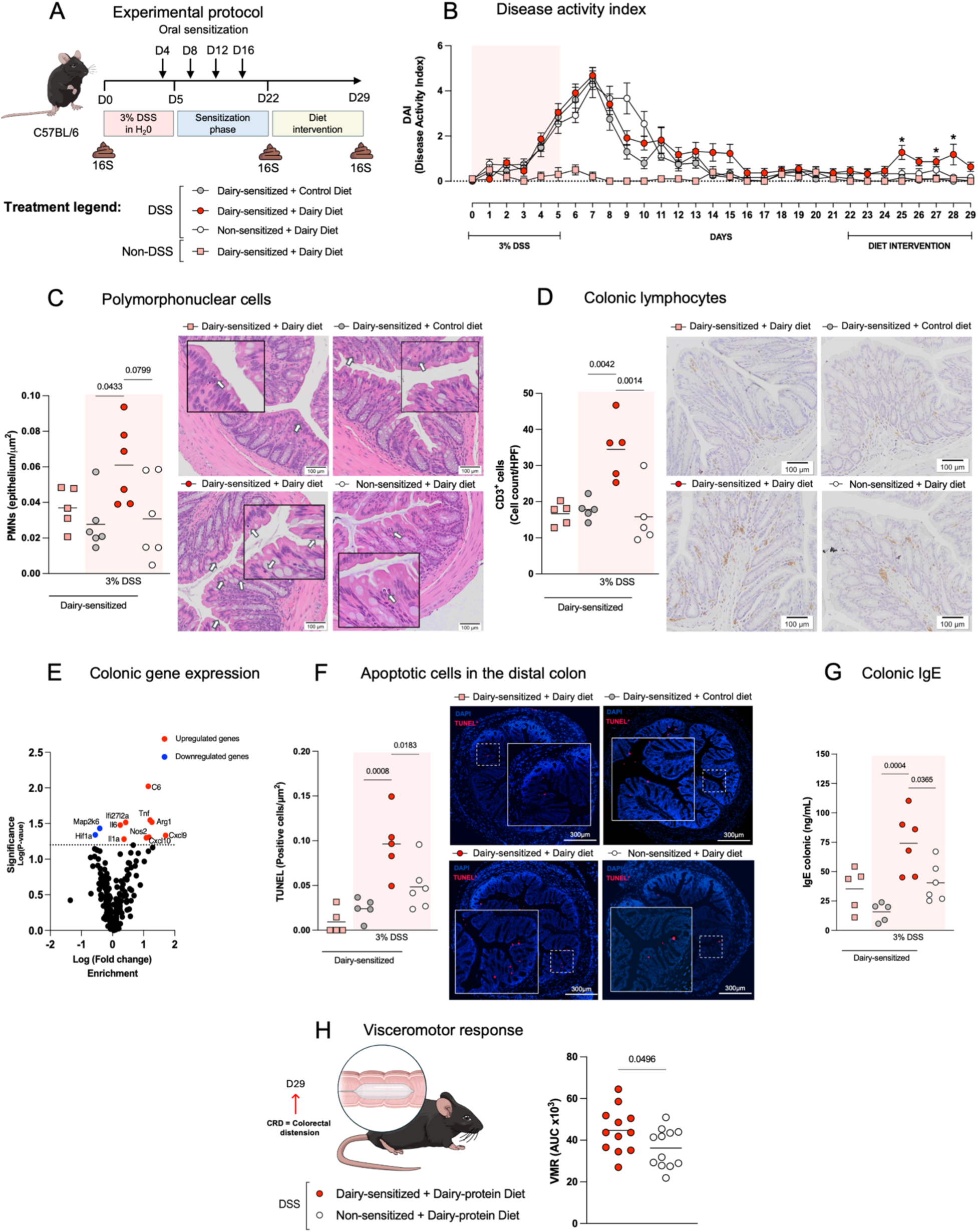
Intestinal inflammation facilitates sensitization to dairy proteins. A. Experimental protocol. Mice were subjected to 3% DSS-induced colitis, followed by oral sensitization to dairy proteins (4x) during the recovery phase, and then a dietary intervention with either a dairy or control diet. Non-DSS controls received water only. B. Disease Activity Index (DAI). C. Colonic Polymorphonuclear cells (PMNs). D. Colonic CD3^+^ cells. E. Colonic gene expression in sensitized mice after DSS-inflammation fed dairy *vs.* control diet. F. Apoptosis cells in distal colon. G. Colonic IgE levels. H. Area under the curve (AUC) of visceromotor responses (VMRs) to colorectal distension (CRD) in sensitized and non-sensitized mice after DSS fed a dairy diet. Data are expressed as mean +/– SEM (B) and mean where each dot represents one mouse (C–D, F–H). Displayed *p* values were calculated using two-way (B) or one-way ANOVA with Bonferroni’s post-hoc test (C-G), and unpaired *t* test with Tukey’s post-hoc test (F). * P <0.05 compared to DSS + non-sensitized + dairy-protein diet group.

To investigate whether inflammation also facilitates sensitization to other dietary antigens, we established a parallel model using gluten, a driver of clinical responses in IBD^3^. C57BL/6 mice were exposed to DSS and sensitized to gluten, then fed either a wheat gluten-containing or control diet (Figure 2A). As in the dairy model, all DSS-treated mice experienced weight loss and increased disease activity, which resolved after recovery (Supplementary Figure 2A and 2B). Upon gluten re-exposure, sensitized mice displayed increased PMNs, CD3^+^ cells, apoptotic cells (Figure 2B-D and Supplementary Figure 2E), and elevated colonic IgE (Figure 2E). Together, these results suggest that intestinal injury promotes sensitization to dairy and gluten proteins, leading to food-dependent immune activation and adverse reactions.

**Figure 2.**
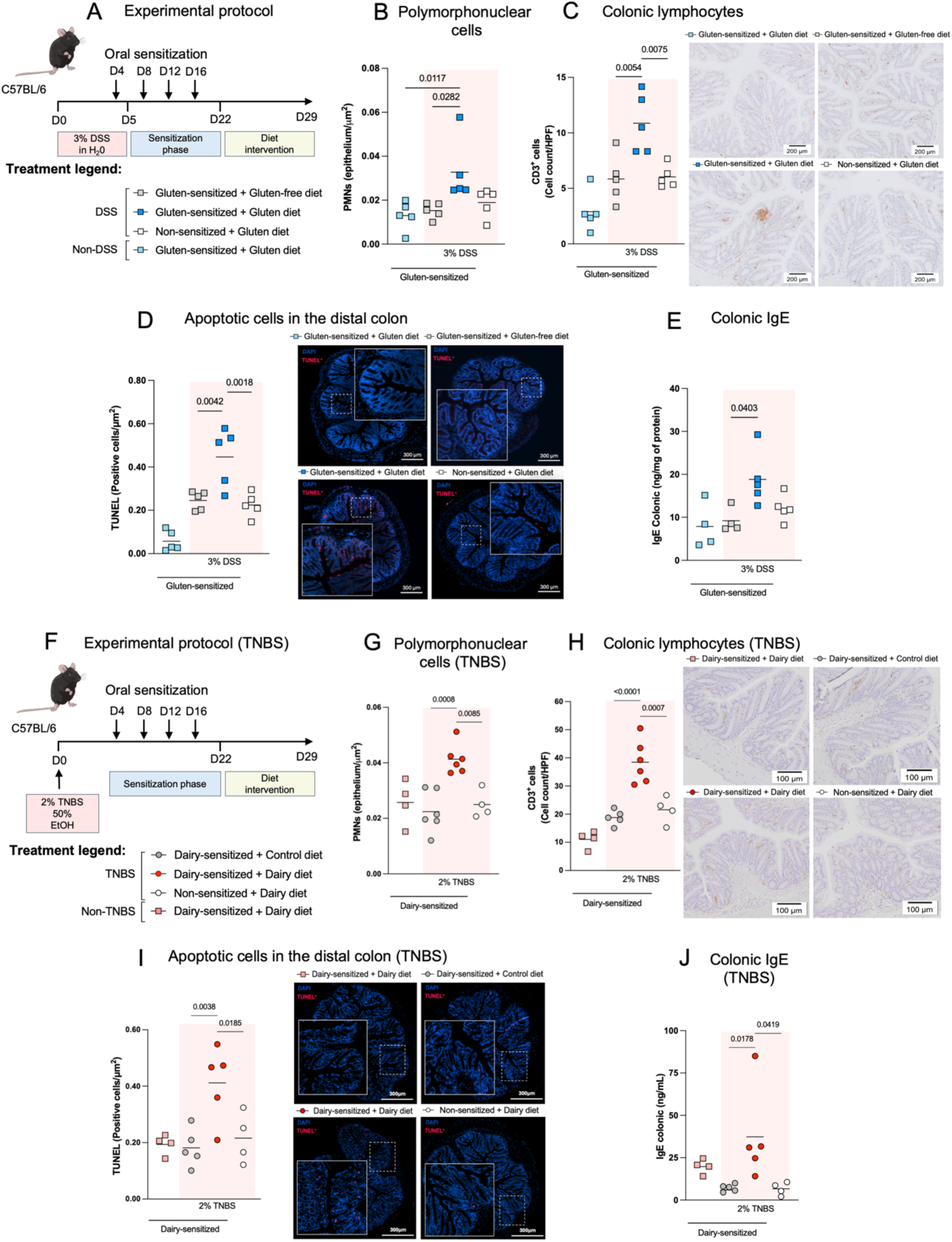
Intestinal inflammation facilitates sensitization to dietary proteins regardless of the type of inflammatory stimulus. A. DSS experimental protocol. Mice were subjected to 3% DSS-induced colitis, followed by oral sensitization to gluten proteins (4x) during the recovery phase, and then a dietary intervention with either a wheat or control diet. Non-DSS controls received water only. B. Polymorphonuclear cells (PMNs) in distal colon. C. CD3^+^ cells in distal colon. D. Apoptotic cells in distal colon. E. Colonic IgE levels. F. TNBS experimental protocol. Mice were subjected to 2% TNBS-induced colitis, followed by oral sensitization to dairy proteins (4x) during the recovery phase, and then a dietary intervention with either a dairy or control diet. Non-TNBS controls received water only. G. PMNs in distal colon. H. CD3^+^ cells in distal colon. I. Apoptotic cells in distal colon. J. Colonic IgE levels. Data are expressed as mean +/– SEM (B), and mean where each dot represents one mouse. Displayed *p* values were calculated using one-way ANOVA (B–J)), with Bonferroni’s post-hoc test.

To study whether food sensitization is facilitated by inflammatory stimuli beyond DSS, we employed a hapten-induced model of colitis using 2,4,6-trinitrobenzene sulfonic acid (TNBS). C57BL/6 mice were exposed to TNBS, sensitized to dairy proteins or sham, and then placed on either a dairy-containing or control diet (Figure 2F). All TNBS-treated mice exhibited weight loss and increased DAI during colitis induction, which resolved after two weeks. During the dietary intervention, however, TNBS-colitis sensitized mice showed significant reductions in body weight and increased DAI compared to controls (Supplementary Figure 2C). Consistent with the DSS model, these mice exhibited increased PMNs, CD3^+^ and apoptotic cells in colonic tissue than non-TNBS and non-sensitized controls (Figure 2G-I and Supplementary Figure 2F), and elevated colonic IgE (Figure 2J), despite mild intestinal inflammation (Supplementary Figure 2D). Altogether, these results suggest that intestinal inflammation, whether induced by DSS or TNBS, promotes sensitization to dietary antigens and drives food-dependent immune responses.

### Mast cells in the colon mediate adverse reactions to food after inflammation

Mast cells have been previously implicated in the development of food-induced adverse reactions^23^. Sensitized-mice previously exposed to inflammation and then fed a dairy- or gluten-diet showed increased tryptase-positive cells in the colonic mucosa compared to non-colitis and non-sensitized controls in both DSS and TNBS-models (Figure 3A and 3B). This suggests mast cell recruitment to the colon in mice sensitized to proteins during active intestinal inflammation. In line with these observations, dairy consumption in sensitized mice post-inflammation increased systemic mucosal mast cell protease-1 (mMCPT-1), a marker of mucosal mast cell degranulation (Figure 3C). Similar results were observed in gluten-sensitized mice after inflammation, with increased tryptase-positive cells and mMCPT-1 levels systemically upon gluten diet consumption (Supplementary Figure 3A and 3B).

**Figure 3.**
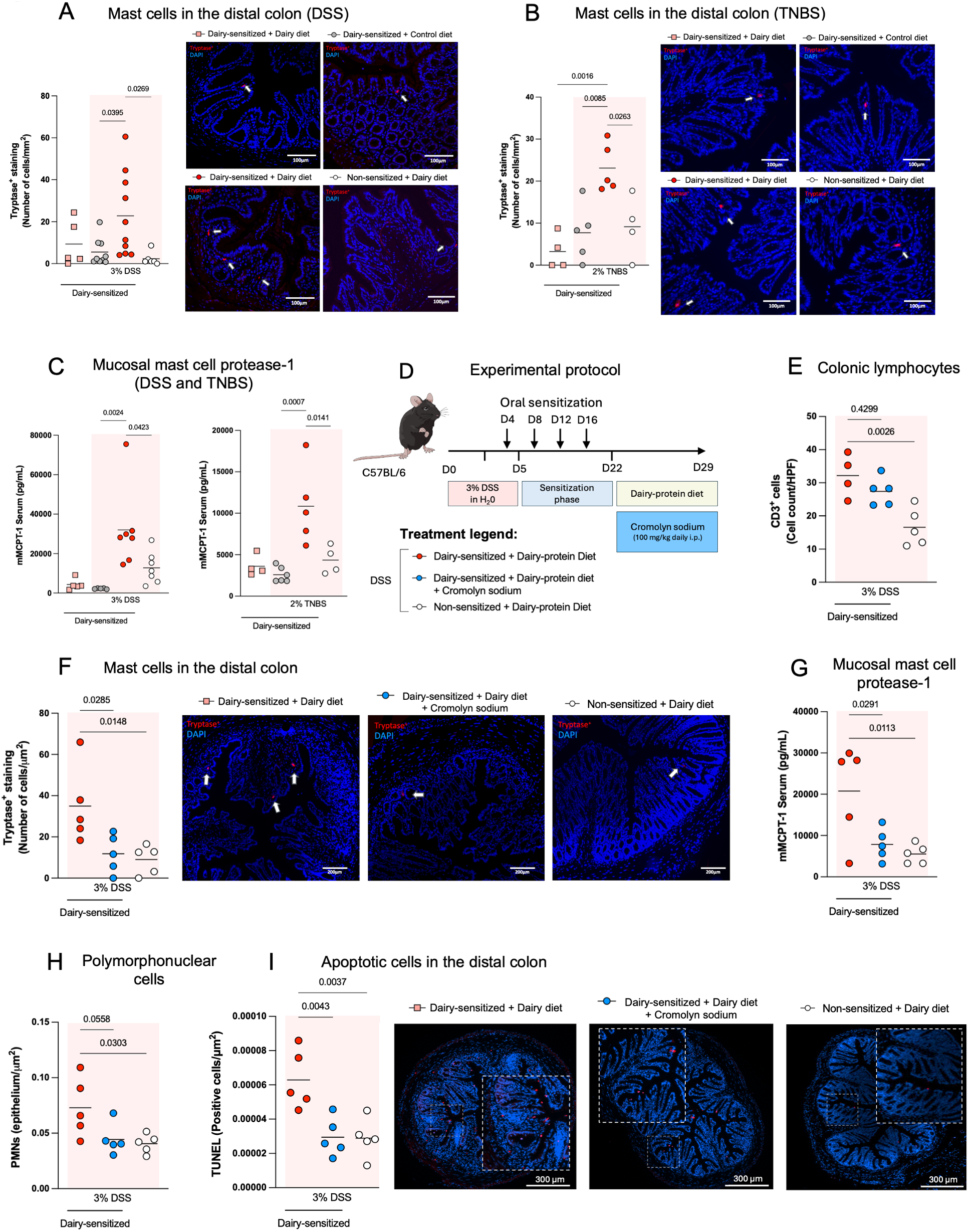
Colonic mast cells mediate adverse food reactions after inflammation. A-B. Tryptase-positive mast cells in the colon and C. Serum mMCPT-1 in mice subjected to DSS and TNBS colitis, sensitization to dairy proteins, and subsequent dietary intervention with either a dairy or control diet. D. Experimental design for mast cell blockade. Mice were subjected to 3% DSS-induced colitis, followed by oral sensitization to dairy proteins (4x) during the recovery phase, and then a dietary intervention with dairy. During dietary intervention, a subset of sensitized mice received daily intraperitoneal cromolyn sodium to inhibit mast cell activation. E. Colonic CD3+ cells. F. Colonic tryptase-positive mast cells. G. Serum mMCPT-1 levels. H. Polymorphonuclear cells in colonic tissue. I. Apoptotic cells in colon. Data are expressed as mean where each dot represents one mouse. Displayed *p* values were calculated using one-way ANOVA with Bonferroni’s post-hoc test.

To investigate the functional involvement of mast cells, C57BL/6 mice were sensitized to dairy proteins or sham post-DSS, and then treated with the mast cell stabilizer cromolyn sodium during dairy consumption (Figure 3D). Mast cell stabilization showed a non-significant DAI reduction during dairy intake (Supplementary Figure 3C) and no changes in colonic CD3^+^ cells (Figure 3E). However, cromolyn sodium significantly reduced both colonic tryptase-positive cells and systemic mMCPT-1 (Figure 3F and 3G). Moreover, cromolyn sodium reduced the number of PMN and apoptotic cells (Figure 3H and 3I), confirming the role of mast cells in food-related immune responses post-inflammation. These results suggest that mast cells are important mediators of food sensitization and food-related immune activation in the colon after inflammation.

### Gut inflammation leads to alterations in microbiota and metabolism of food triggers

To study the involvement of the microbiota in inflammation-driven sensitization, we analyzed the fecal microbiota of DSS-exposed mice sensitized to dairy proteins (Figure 1A). DSS led to profound changes in the fecal microbiota, characterized by reduction in alpha diversity and different beta diversity clustering (Figure 4A and 4B and Supplementary Figure 4A and 4B). Core colonic taxa with known food degrading capacity^16^, including *Bifidobacterium* and *Clostridium,* were notably depleted (Figure 4C and 4D and Supplementary Figure 5A). To assess the functional implications of these alterations, we used *Bifidobacterium* and *Clostridium* strains previously isolated from feces of healthy humans and incubated them with dairy proteins *in vitro*. *Bifidobacterium* and *Clostridium* strains efficiently degrade β-casein (Bos d 11) and β-lactoglobulin (Bos d 5), the main antigenic components in dairy (Figure 4E), suggesting that inflammation may disrupt microbial populations critical for dietary antigen processing.

**Figure 4.**
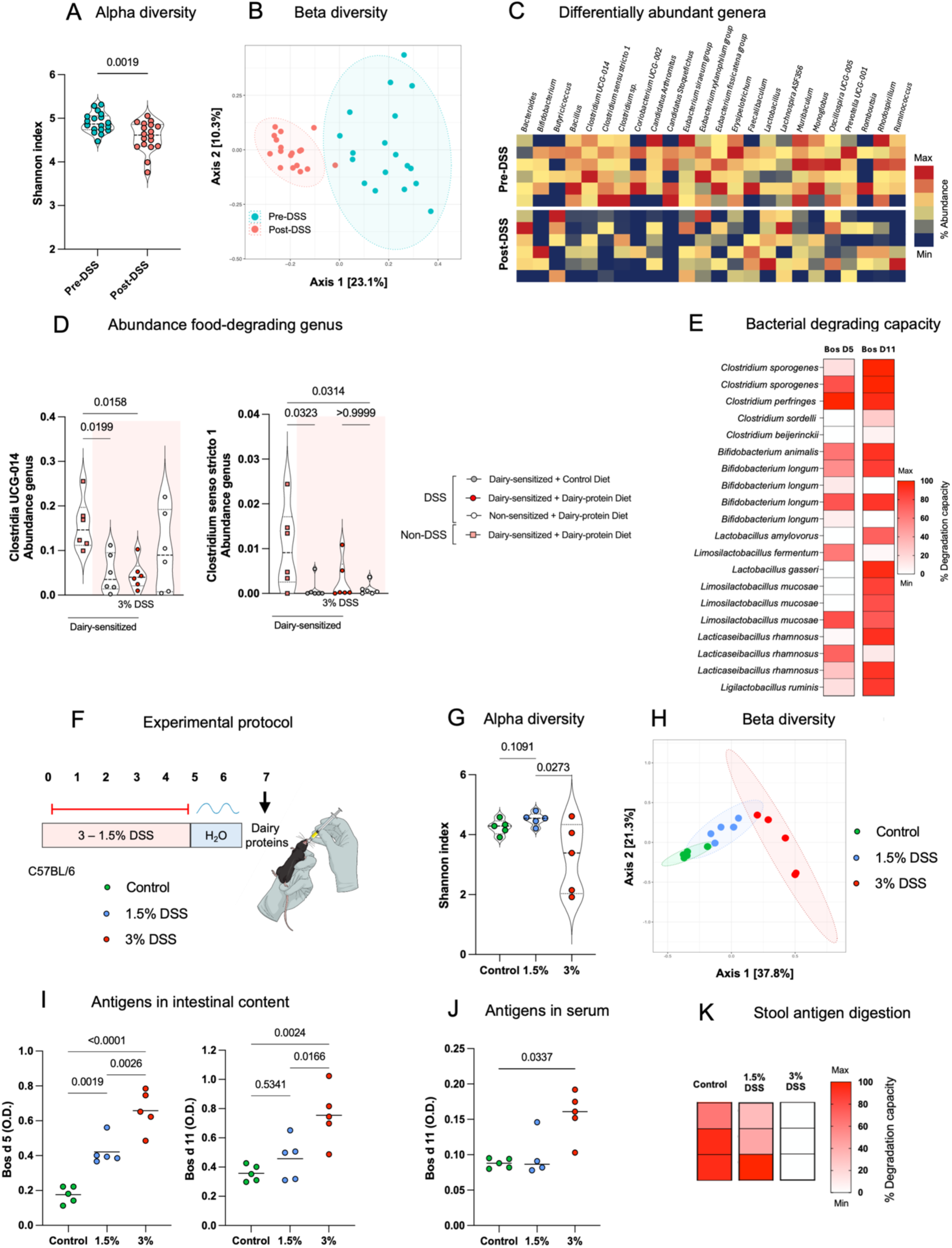
Gut inflammation alters gut microbiota and food antigen metabolism. A. Alpha-diversity (Shannon index) and B. Beta-diversity (Bray–Curtis dissimilarity) of colonic microbiota of dairy- and sham-sensitized mice before and after DSS treatment, prior to dietary intervention. C. Heatmap of differentially abundant taxa at pre-DSS, post-DSS, (blue=0%, red=100% relative abundance). D. Relative abundance of two *Clostridiales* genera in dairy sensitized mice after DSS inflammation fed either a dairy or control diet. E. Heatmap of bacterial degradation capacity for β-lactoglobulin (Bos d 5) and β-casein (Bos d 11) using human bacterial isolates (white=0%, red=100% degradation relative to dairy antigens). F. Experimental protocol for microbial dairy metabolism after inflammation. Mice were subjected to 0% (control), 1.5%, or 3% DSS-induced colitis, followed by oral gavage with the dairy proteins β-casein and β-lactoglobulin. G. Alpha-diversity (Shannon index). H. Beta-diversity (Bray-Curtis dissimilarity) of colonic microbiota. (I) Intestinal and (J) systemic levels of Bos d 5 and Bos d 11. K. Heatmap of the average stool antigen digestion capacity (blue=0%, red=100% degradation relative to dairy antigens). Data are shown as violin plots where each dot represents one mouse and median values are shown (A, C, G), as mean where each dot represents one mouse (I, J). Statistical analyses were performed using Kruskal-Wallis test with Dunn’s test post hoc test, or one-way ANOVA with Bonferroni’s post-hoc test.

To validate these findings, we administered escalating DSS doses to C57BL/6 mice followed by oral dairy protein challenge (Figure 4F). DSS led to dose-dependent microbiota alterations (Figure 4G and 4H), with pronounced reduction of dairy-metabolizing taxa like *Clostridium* at higher concentrations (Figure 4G-H, Supplementary Figure 4C). Systemic and gastrointestinal quantification of dairy antigens revealed elevated Bos d 5 and Bos d 11 levels in intestinal luminal contents and serum of DSS-treated mice (Figure 4I and 4J). Feces from inflamed mice also exhibited reduced *ex vivo* dairy antigen-degrading capacity against Bos D 11 (Figure 4K). While DSS led to increased intestinal damage and PMN cell infiltration, no differences in colonic IgE or mMCPT-1 levels were observed, confirming that inflammation is not sufficient to promote food-driven immunity (Supplementary Figure 5A-D). Collectively, the data indicate that gut inflammation disrupts commensal taxa responsible for dietary antigen breakdown, impairing local metabolic clearance of food antigens. This dysfunction may permit increased systemic exposure to immunogenic peptides, a critical step of food sensitization, potentially linking mucosal inflammation to adverse food responses.

### Altered intestinal microbiota by inflammation facilitates sensitization to food

We next evaluated the involvement of the microbiota in food sensitization using germ-free C57BL/6 mice colonized with SPF microbiota (non-DSS microbiota) or post-inflammatory SPF microbiota induced by DSS (post-DSS microbiota). Mice were then sensitized to dairy proteins using cholera toxin and placed on dairy-containing diets (Figure 5A). Mice with post-DSS microbiota exhibited increased DAI after sensitization when fed dairy diet, a response absent in mice with non-DSS microbiota (Figure 5B). Although no differences were observed in overt inflammation between groups (Supplementary Figure 6A), mice with post-DSS microbiota presented increased colonic PMN infiltration and CD3^+^ lymphocyte counts compared to non-DSS microbiota recipients (Figure 5C and 5D). Furthermore, post-DSS microbiota was associated with elevated colonic IgE levels (Figure 5E), increased systemic mMCPT-1 (Figure 5F), and higher numbers of colonic tryptase positive mast cells (Figure 5G). Collectively, these results suggest that inflammation-induced microbial alterations promote food sensitization and adverse reactions to foods.

**Figure 5.**
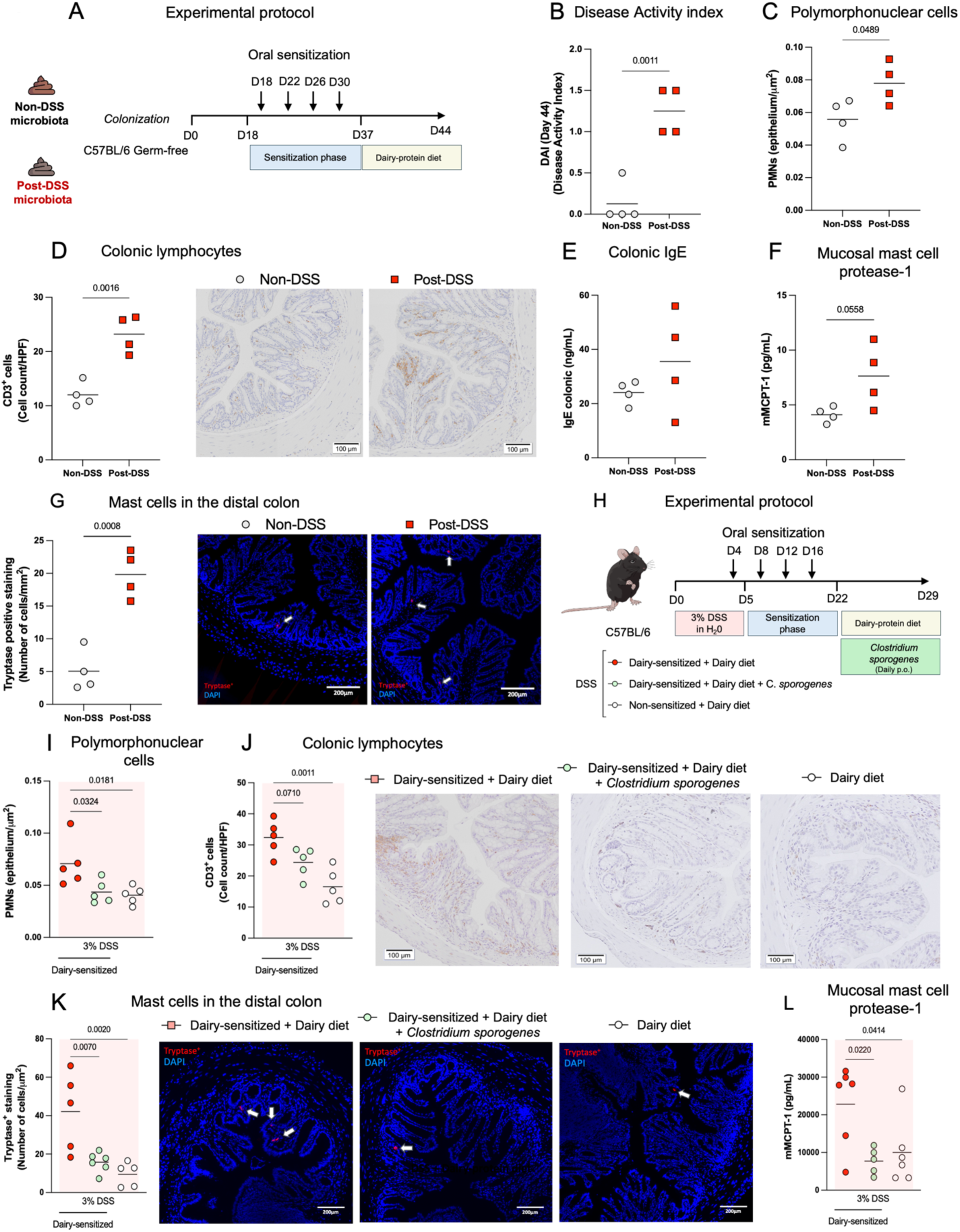
Post-DSS microbiota facilitate sensitization to food antigens and dairy-degrading bacteria mitigate inflammation-driven adverse dairy reactions. A. Microbiota transfer protocol. Germ-free mice were colonized with either non-DSS SPF microbiota or DSS SPF microbiota, followed by oral sensitization (4x) to dairy proteins, and subsequent dietary intervention with dairy. B. Disease Activity Index (DAI). C. Colonic Polymorphonuclear cells (PMNs). D. Colonic CD3^+^ cells. E. IgE serum levels. F. mMCPT-1 serum levels. G. Tryptase-positive mast cells in colon. H. Bacterial supplementation protocol. Mice were exposed to DSS-induced colitis followed by oral sensitization (4x) to dairy proteins. Mice were subsequently challenged with dairy-protein diet and supplemented daily with antigen-degrading *Clostridium sporogenes* or sham. I. Colonic polymorphonuclear cells. J. Colonic CD3+ cells. K. Colonic tryptase-positive mast cells. L. mMCPT-1 serum levels. Data are shown as mean where each dot represents one mouse. Statistical differences were calculated using unpaired t test (B-G) or one-way ANOVA with Bonferroni’s post-hoc test (I-L).

### Dairy-degrading bacteria improve adverse reactions to foods associated to inflammation

Given the microbiota’s capacity to digest food triggers, we investigated whether supplementation with antigen-degrading bacteria could mitigate inflammation-associated adverse food reactions. C57BL/6 mice were exposed to DSS, sensitized to dairy proteins, and subsequently fed a dairy-containing diet with or without supplementation of *Clostridium sporogenes*, a dairy antigen-degrading bacterium that is depleted during inflammation (Figure 5H). Bacterial supplementation did not significantly improve DAI (Supplementary Figure 7A), but it reduced PMN infiltration in the colonic mucosa (Figure 5I). While a modest reduction in CD3^+^ cells was observed in the colonic mucosa (p=0.07; Figure 5J), *C. sporogenes* supplementation decreased tryptase-positive mast cell accumulation in the distal colon (Figure K) and reduced systemic levels of mMCPT-1 (Figure 5L). These findings highlight the importance of microbial antigen metabolism in regulating inflammation-associated adverse food reactions and emphasize the potential therapeutic relevance of microbial interventions.

### Dairy exacerbates colitis in sensitized mice after inflammation via microbial metabolism

IBD patients frequently report that food triggers worsen inflammation during disease flares^24,25^. To investigate this phenomenon, we examined whether sensitization to dietary antigens following inflammation affects colitis severity upon re-exposure to the triggering food. C57BL/6 mice were sensitized to dairy or sham after DSS-induced inflammation and subsequently exposed to a second cycle of DSS while consuming either dairy-containing or control diets (Figure 6A). Dairy-sensitized mice fed the dairy diet exhibited significantly increased DAI, reduced colon length, and more severe histologic damage compared to non-sensitized controls and sensitized mice fed controls diets (Figure 6B–D). In addition, these mice also displayed increased numbers of tryptase-positive mast cells and increased epithelial apoptosis in colonic tissue (Figure 6E and 6F), suggesting that re-introduction of food triggers exacerbates colitis via mast cell activation and enhanced epithelial injury.

**Figure 6.**
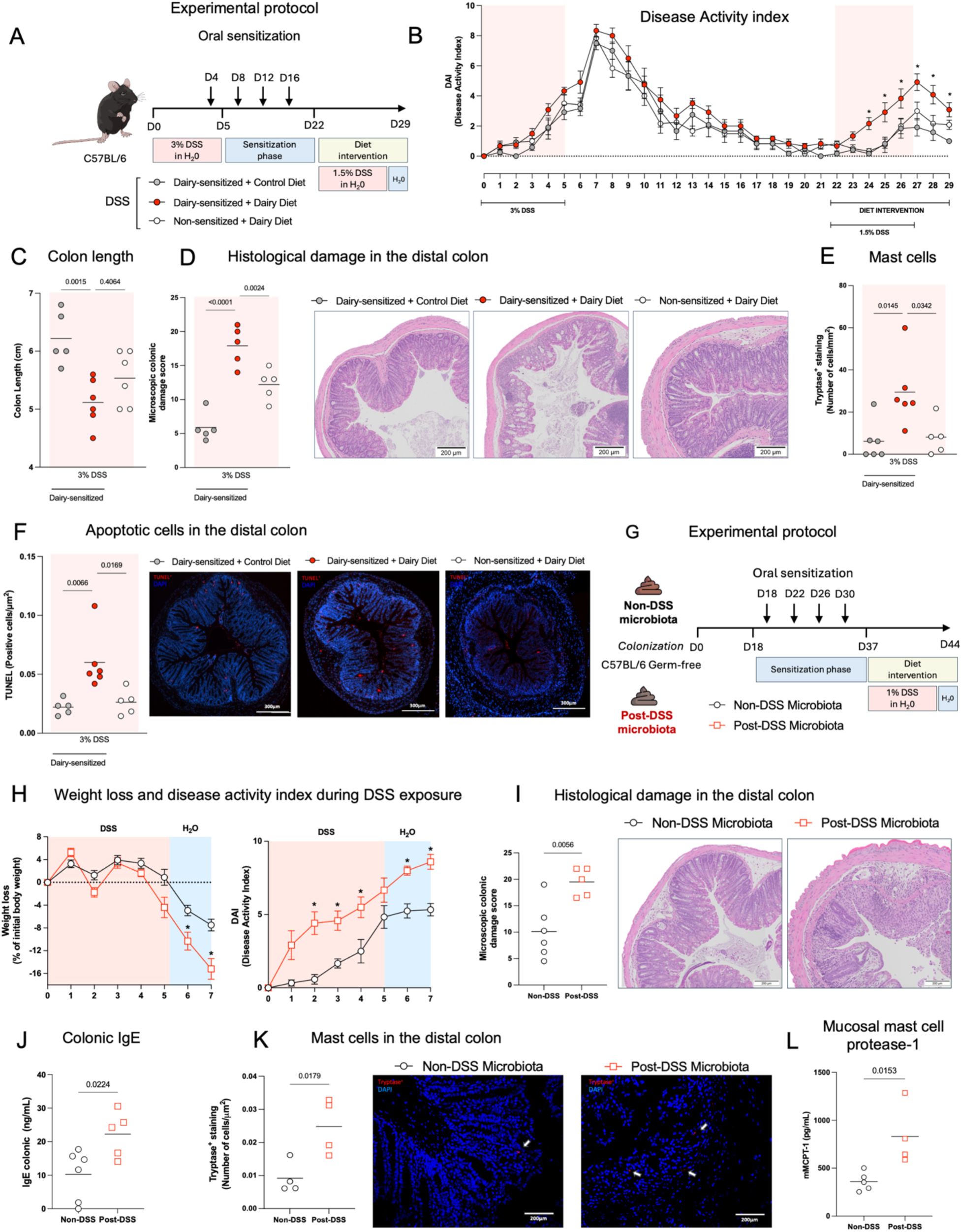
Dairy exacerbates colitis in sensitized mice after inflammation via microbial metabolism. A. Experimental colitis protocol. Mice were subjected to 3% DSS-induced colitis, followed by oral sensitization to dairy proteins (4x) or sham during the recovery phase, and then a dietary intervention with either a dairy or control diet. Mice were subjected to a second mild 1.5% DSS challenge during the dietary intervention. B. Disease activity index. C. Colon length at endpoint. D. Microscopic damage score of distal colons. E. Tryptase-positive mast cells in distal colon. F. TUNEL+ staining of apoptotic cells in distal colon. G. Microbiota transfer protocol. Germ-free mice were colonized with either wild-type non-DSS SPF microbiota or post-DSS SPF microbiota, followed by oral sensitization (4x) to dairy proteins, and subsequent dietary intervention with dairy. Mice were subjected to a second mild 1.0% DSS challenge during the dietary intervention. H. Body weight change (left panel) and disease activity index (right panel) during DSS. I. Microscopic damage score of distal colon. J. Colonic tissue IgE levels. K. Tryptase-positive mast cells in distal colon. L. Serum mMCPT-1 levels. Data are shown as mean +/− SEM (B, H) or mean where each dot represents one mouse (C, D, E, F, I, J, K. L). Statistical analyses were performed with one-way ANOVA or two-way ANOVA with Bonferroni’s post-hoc test, and unpaired t test (I-L). * P <0.05 compared to the non-sensitized group receiving dairy-protein diet.

To study whether food-driven colitis worsening was mediated by inflammation-induced microbial alterations, germ-free C57BL/6 mice were colonized with either SPF (Non-DSS) or post-DSS microbiota as described above. Following colonization, mice were sensitized to dairy proteins and subsequently challenged with DSS while consuming a dairy diet (Figure 6G). Mice with post-DSS microbiota developed significantly greater weight loss and disease activity during DSS exposure compared to those with wild-type microbiota (Figure 6H). Histologic assessment revealed more severe mucosal damage in mice colonized with post-DSS microbiota (Figure 6I). Moreover, mice with post-DSS microbiota showed elevated IgE levels in colonic mucosa (Figure 6J), increased tryptase-positive mast cells in distal colon (Figure 6K) and increased serum levels of mMCPT-1 (Figure 6L), compared to mice with non-DSS microbiota. To confirm that colitis worsening was not driven by DSS-associated microbiota independently of food sensitization, we repeated the experiment without sensitizing mice to dairy proteins. No differences in weight loss, disease activity index (DAI), or colonic IgE levels were observed between mice colonized with microbiota from non-DSS or post-DSS donors. Only a minor increase in colonic PMN infiltration was observed in mice colonized with post-DSS microbiota (Supplementary Figure 8A–E). These findings demonstrate that inflammation-induced microbial alterations promote heightened sensitivity to dietary antigens and exacerbate subsequent inflammatory responses, establishing a link between microbiota alterations, food sensitization, and colitis severity.

### Self-reported adverse food reactions in IBD associate with alterations in fecal microbiota and depletion of antigen-degrading bacteria

To translate our experimental findings to human disease, we investigated the relationship between food intolerances, IBD diagnosis, and gut microbiota composition in a clinical cohort. IBD patients in remission (N=104; 58 with CD and 46 with UC) and non-IBD controls (N=44) were recruited and surveyed on adverse food reactions and food-driven symptomatology (Figure 7A, Supplementary Figure 9A and Supplementary Table 1A). Food intolerance was more common in IBD patients (85.3% CD, 86.2% UC) than in non-IBD controls (33.3%) (Figure 7B), and involved a greater number of foods: CD averaged 3.5 (SD = 1.90), UC 3.2 (SD = 1.80), versus 1.3 (SD = 0.83) in non-IBD (Supplementary Figure 9B). The most reported food triggers in IBD included dairy, wheat, peanuts/tree nuts, caffeine and fiber (Figure 7C). IBD patients reported discomfort, bloating, diarrhea, and abdominal pain upon their consumption (Supplementary Figure 9C). Additionally, about 40% of IBD patients reported food-associated symptoms prior to diagnosis, while 60% reported a worsening of food-related symptoms following diagnosis. Interestingly, 76% of patients with UC reported the onset of food intolerances after their clinical diagnosis of IBD, suggesting that intestinal inflammation may play an important role in the development of adverse food reactions (Figure 9D and Supplementary Figure 9D).

**Figure 7:**
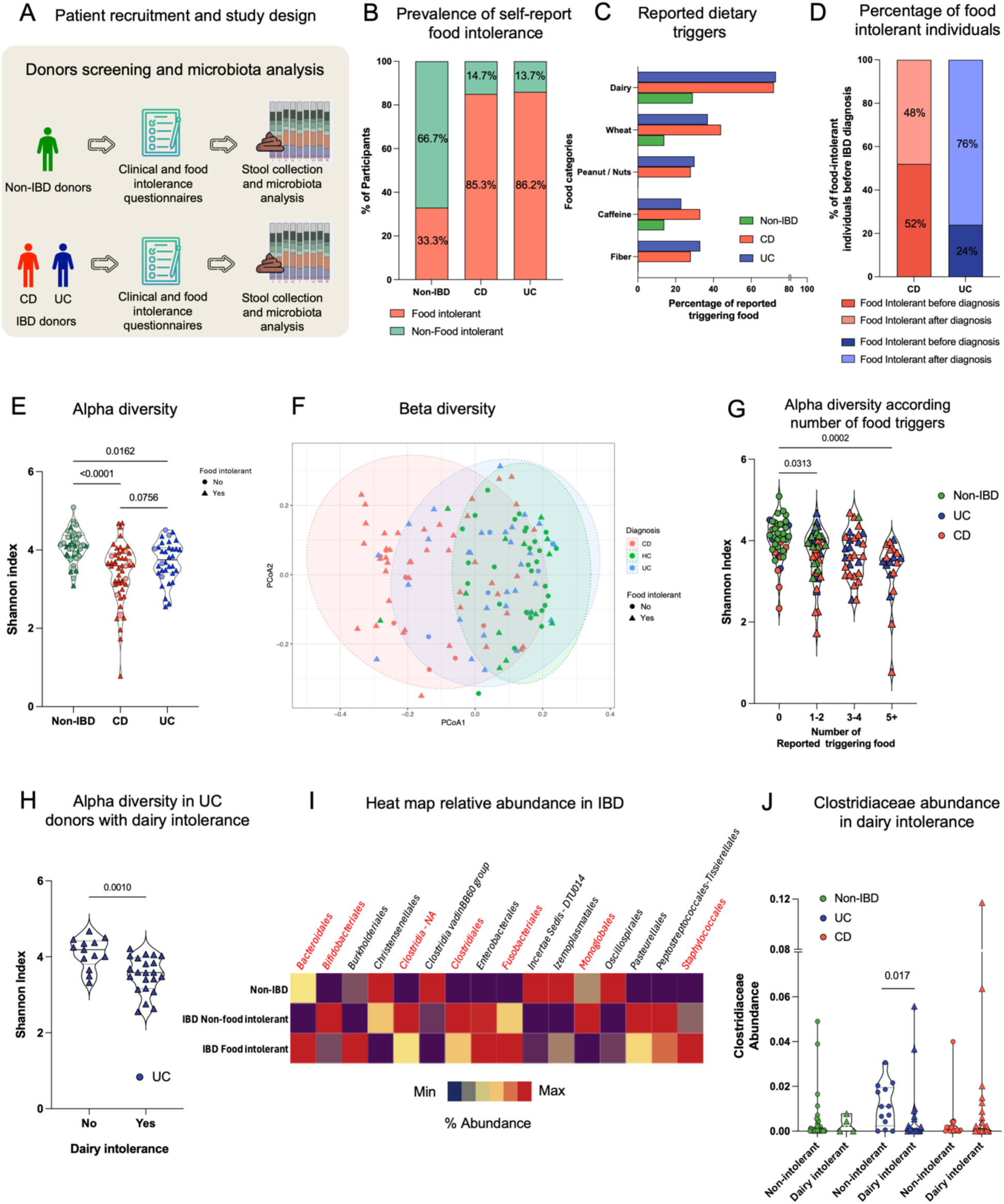
IBD patients report increased food intolerances and exhibit altered fecal microbiota compared to healthy controls. A. Study design. 104 IBD patients (58 CD and 46 UC), and 44 non-IBD controls were recruited. Each study participant responded to clinical, and food related adverse reaction questionnaires and provided stool samples for microbiota analysis. B. Percentage of participants self-reporting adverse food reactions across CD (85.3%), UC (86.2%), and non-IBD (33.3%). C. Distribution of common food triggers reported by participants with self-reported food intolerances, categorized by food type. D. Percentage of patients reporting food intolerance starting before and after IBD diagnosis. E. Alpha diversity (Shannon index) and F. Beta diversity plot (Bray Curtis) in UC (Blue), CD (Red), and non-IBD participants (Green), with reported adverse food reactions (shape; triangle: Yes, Circle: No). G. Shannon alpha diversity based on the number of self-reported food intolerances across participant groups. H. Alpha diversity in UC patients with and without self-reported intolerance to dairy. I. Heatmap of bacterial order-level relative abundance in non-IBD controls and IBD patients with and without self-reported food intolerances; Heat map; blue=0%, red=100% relative abundance; red-highlighted taxa denote statistically significant differences in relative abundance between IBD patients with vs. without food intolerance. J. Relative abundance of *Clostridiaceae* in participants with and without self-reporting dairy intolerance, stratified by diagnostic group. Statistical significance was calculated using Kruskal Wallis with Dunn’s post-hoc test. Same as above for stats and each dot represents one donor.

Fecal microbiota analysis (Supplementary Table 2) revealed that disease, rather than food intolerance, was the primary driver of microbiota variation. IBD patients exhibited significantly reduced alpha diversity and distinct beta diversity clustering patterns compared to non-IBD controls (Figure 7E-F). Of interest, reduced alpha diversity associated with the number of food intolerances in both non-IBD and UC patients (Figure 7G). Curiously, reduced alpha diversity was also associated with adverse food reactions in the absence of IBD (Supplementary Figure 9E.) Moreover, UC patients reporting dairy intolerance displayed significantly lower alpha diversity compared to UC patients without dairy-related symptoms (Figure 7H). IBD patients reporting food intolerances also exhibited depletion of specific bacterial taxa, particularly members of the *Clostridiaceae* family involved in the metabolism of food triggers (Figure 7I). Most notably, UC patients with dairy intolerance showed significant reductions in *Clostridium* sensu stricto 1 and Clostridia UCG-14 (Figure 7J). Altogether, these findings establish a clinical association between IBD, adverse food reactions, and altered gut microbiota composition, particularly reduction of microbial complexity and depletion of bacterial taxa capable of metabolizing food triggers.

## Discussion

Approximately 20% of the population reports adverse reactions to foods. While certain food sensitivities, such as allergies and celiac disease (CeD), are well-defined, the causes behind other self-reported intolerances remain less clear ^12^. This is particularly true for individuals with chronic disorders such as irritable bowel syndrome and IBD. Our findings show that a significant majority of IBD patients report adverse food reactions during remission and acknowledge that certain foods exacerbate inflammation and symptoms. In this study, we demonstrate that intestinal inflammation promotes sensitization to food proteins by disrupting the microbial metabolism of food triggers (e.g. dairy and gluten) and eliciting mast cell-mediated immune responses. Re-exposure to these foods caused visceral pain and immune activation in sensitized mice and further exacerbated inflammation during experimentally induced colitis. We also confirmed that bacteria involved in the metabolism of food triggers such as dairy or gluten proteins are reduced in IBD patients who self-report food intolerances.

IBD patients may continue to experience persistent gastrointestinal symptoms even after successful clinical and histological remission. Research shows that 18–24% of IBD patients report symptoms like abdominal pain, changes in bowel habits, bloating, and increased gas despite having no clear signs of active inflammation^26^. These ongoing GI symptoms are linked to psychological issues, decreased quality of life, and higher healthcare use^27^. Both patients and clinicians acknowledge the role of diet in managing IBD, with patients identifying certain foods as triggers for their symptoms, prompting them to follow elimination diets. Unfortunately, these diets can lead to nutritional deficiencies and other health problems, and targeted adjuvant therapies are needed. Our study shows that more than 80% of IBD patients, regardless of their condition, self-reported food intolerances, a finding consistent with prior research^3^. Despite the concerning prevalence of self-reported food intolerances in IBD, the primary dietary triggers and the underlying mechanisms remain poorly understood.

In both our study and other published research, the most commonly self-reported food reactions in IBD are to dairy and wheat^28^. Since these are primary triggers for lactose intolerance and CeD respectively, many of these patients are frequently labeled with these conditions despite a lack of diagnostic confirmation. However, non-canonical immune responses to components of wheat and dairy have been shown in patients with functional intestinal disorders. These include barrier dysfunction, immune activation (CD3+ cells), and eosinophil activation^29,30^. Indeed, local immune responses to gluten and dairy drive meal-induced abdominal pain in patients with functional disorders via IgE^+^ mast-cells^23^. In the same study, the authors show that microbial infections can trigger immune responses that lead to the production of dietary-antigen-specific IgE antibodies in mice. Following re-exposure to the food antigen, an IgE- and mast-cell-dependent mechanism triggered an increase in visceral pain^23^. While confirmation of these responses in the mucosa of IBD patients is still pending, our preclinical results indicate a similar phenomenon after inflammation. Inflammatory stimuli (chemically- and hapten-induced) promote food sensitization and the reintroduction of the food triggers leads to immune activation and visceral hypersensitivity. These responses are characterized by increased CD3^+^ cells and apoptosis in the colon through IgE and mast-cell dependent mechanisms. Of note, mast cells have been linked to epithelial apoptosis through their chymase enzyme via TNF-alpha-mediated pathways^31^, and upregulation of TNF-alpha was observed in our study (Figure 1E). In addition, IBD patients exhibit elevated levels of circulating IgE^+^ or FcεRIα^+^ immune cells, similar to allergic patients^32^, and antibodies to cow’s milk proteins^33^. Together, our findings and those from previous studies propose a novel IgE-mediated mechanism of food sensitivity, driven by inflammation or other microbial triggers, which may explain adverse food reactions in patients with chronic intestinal disorders.

Inflammation is characterized by increased intestinal permeability and robust non-specific immune reactions, both of which can help promote food sensitization. It is noteworthy that 76% of UC patients reported the onset of food intolerances after their clinical diagnosis of IBD, indicating that inflammation may be a key factor driving food intolerance. Furthermore, IBD patients exhibit changes in microbial composition and function ^14,15^, and microbes are important drivers of food sensitization^13^. Indeed, several mechanisms through which microbes contribute to the development of food sensitivity have been identified^34^. Among these, the ability of the microbiota to effectively metabolize dietary antigens is crucial^10,11,35^, as most of these antigens are resistant to full digestion by human digestive enzymes^36^. In this study, we show that inflammation causes significant changes in microbiota composition and its ability to digest food triggers in preclinical models. Indeed, microbial members such as *Clostridium* with the capacity to digest dairy and gluten proteins are depleted after inflammation. We also confirmed that microbial changes following inflammation drive food sensitization and food-induced adverse reactions in cross-microbiota transfer experiments using gnotobiotic models. As previously reported in the literature^37^, we also observed changes in the microbiota of IBD patients, including reduced alpha-diversity and a depletion of core commensals. Interestingly, alpha diversity was linked to the total number of self-reported food intolerances, and particularly to dairy intolerance. IBD patients who reported food intolerances also showed a depletion of taxa, including well-known bacterial species involved in the metabolism of food triggers, such as members of *Clostridium*. Overall, our findings suggest that microbiota disruptions in IBD play a key role in driving adverse reactions to foods.

The main treatment for patients with well defined food sensitivities, such as CeD and food allergy, is the long-life removal of the food trigger. While food-related adverse reactions in IBD are not well understood, our study suggests that IBD patients often adopt various dietary interventions to manage their symptoms, including eliminating foods they identify as problematic. Identifying local mucosal IgE-mediated immune responses to specific antigens, as described in this study, could help stratify food intolerances and tailor individual dietary treatments for IBD patients, thereby improving symptom management and preventing potential nutritional imbalances. Our results also suggest that microbes could provide potential benefits in managing food-related adverse reactions in IBD. We demonstrated that supplementing with antigen-degrading bacteria, which are depleted during inflammation, helps reduce immune and clinical responses triggered by foods. These findings support the use of probiotic-like bacteria with dairy or gluten-degrading capabilities in IBD patients with well-characterized food intolerances, emphasizing the therapeutic potential of our study. Further human studies are needed using such probiotics to determine whether they can improve clinical symptoms for IBD patients with food intolerances.

## Abbreviations

AGU: Axenic gnotobiotic unit
Arg-1: Arginase 1
AUC: Area under the curve
BSA: Bovine serum albumin
C6: Complement component 6
CD: Crohn’s disease
CFU: Colony-forming unit
CRD: Colorectal distension
Cxcl10: C-X-C Motif Chemokine Ligand 10
Cxcl9: C-X-C Motif Chemokine Ligand 9
DAB: 3,3’-diaminobenzidine
DAI: Disease activity index
DSS: Dextran sulfate sodium
HPF: High-power field
IBD: Inflammatory bowel disease
IgE: Immunoglobulin E
Il-10: Interleukin-10
Il-6: Interleukin-6
mMCPT-1: Mucosal mouse mast cell protease-1
PBS: Phosphate buffered saline
PMNs: Polymorphonuclear cells
SPF: Specific pathogen free
Th1: T-helper type 1 cells
TNBS: trinitrobenzene sulfonic acid
Tnf: Tumor necrosis factor
UC: Ulcerative colitis
VMR: Visceromotor responses

## CRediT Authorship Contributions

Bruna B. Da Luz, PhD (Writing – review and editing, Writing – original draft, Data curation, Methodology, Investigation, and Formal analysis)

Liam E. Rondeau, PhD (Writing – review and editing, Investigation)

Rebecca Dang, MSc (Data curation, Investigation, Formal analysis)

Dana Coppens, BSc (Investigation, Formal analysis)

Dominika Boron, MSc (Data curation, Investigation)

Pranshu Muppidi BSc (Investigation)

Jessica Linton, BSc (Data curation, Investigation

Fernando A. Vicentini, PhD (Investigation, Formal analysis)

John K. Marshall, MD (Data curation, methodology)

Giada De Palma, PhD (Writing – review and editing, Methodology)

Premysl Bercik, MD, PhD (Writing – review and editing, Methodology)

Neeraj Narula, MD (Conceptualization, Writing – review and editing, Data curation, Methodology, Funding acquisition)

Alberto Caminero, PhD (Conceptualization, Writing – original draft, Writing – review and editing, Data curation, Formal analysis, Investigation, Methodology, Supervision, Funding acquisition)

## Acknowledgments

The authors thanks the Farncombe Family Axenic-Gnotobiotic Facility and McMaster Genomics Facility. Illustrations were created with Mindthegraph.com.

## Data Transparency Statement

Anonymized microbiota data from individual participants, and animal experimental are deposited at GenBank (NIH - PRJNA1301547)

